# Lineage, Identity, and Fate of Distinct Progenitor Populations in the Embryonic Olfactory Epithelium

**DOI:** 10.1101/2021.10.29.466513

**Authors:** Elizabeth M. Paronett, Corey A. Bryan, Thomas M. Maynard, Anthony-S. LaMantia

**Affiliations:** Center for Neurobiology Research, The Fralin Biomedical Research Institute, Virginia Tech-Carilion School of Medicine, Roanoke VA, 24015; Department of Biological Sciences, Virginia Tech, Blacksburg VA, 24061

## Abstract

We defined a temporal dimension of precursor diversity and lineage in the developing mouse olfactory epithelium (OE) at mid-gestation that results in genesis of distinct cell classes. Slow, symmetrically dividing Meis1+/ Pax7+ progenitors in the early differentiating lateral OE give rise to small numbers of Ascl1+ precursors in the dorsolateral and ventromedial OE. Few of the initial progeny of the Ascl1+ precursors immediately generate olfactory receptor neurons (ORNs). Instead, most early progeny of this temporally defined precursor cohort, labeled via temporally discreet tamoxifen-dependent *Ascl1*^Cre^-driven recombination, populate a dorsomedial OE domain comprised of proliferative Ascl1+ as well as Ascl1-cells from which newly generated ORNs are mostly excluded. The most prominent early progeny of these Ascl1+ OE precursors are migratory mass cells associated with the nascent olfactory nerve (ON) in the frontonasal mesenchyme. These temporal, regional and lineage distinctions are matched by differences in proliferative capacity and modes of division in isolated, molecularly distinct lateral versus medial OE precursors. By late gestation, the progeny of the temporally and spatially defined Ascl1+ precursor cohort include few proliferating precursors. Instead, these cells generate a substantial subset of OE sustentacular cells, spatially restricted ORNs, and ensheathing cells associated with actively growing as well as mature ON axons. Accordingly, from the earliest stages of OE differentiation, distinct temporal and spatial precursor identities provide a template for acquisition of subsequent OE and ON cellular diversity.

## INTRODUCTION

The initial specification and subsequent diversification of ectodermal placode-derived stem cells that generate the olfactory sensory epithelium (OE) sestablishes three essential facets of olfactory pathway differentiation and maintenance: precursors that generate olfactory receptor neurons (ORN) throughout life (Balmer and LaMantia, 2005; Sokpor et al., 2018); cellular diversity, including olfactory ensheathing cells that facilitate OE homeostasis and ORN axon growth (Franssen et al., 2007; Rigby et al., 2020), and “zonal” odorant receptor expression for ORN functional diversity (Coleman et al., 2019; Monahan and Lomvardas, 2015; Rodriguez, 2013; Zapiec and Mombaerts, 2020). Developing and mature OE precursors have been classified based upon transcription factor expression; however, OE position, signaling, or lineage dynamics also distinguish these cells (Kawauchi et al., 2005; LaMantia et al., 2000; Shou et al., 2000; Whitesides et al., 1998). It remains uncertain, however, whether additional distinctions contribute to OE cellular diversity. We asked whether OE precursors (Beites et al., 2005; Kam et al., 2014; Tucker et al., 2010) are divided into temporally diverse subsets with distinct positions and fates during the earliest stages of OE patterning and differentiation.

Developing and mature OE precursor molecular diversity has been explored in detail (reviewed by Schwob et al., 2017). Multiple transcription factors have been associated with these cells, including Sox2 (Donner et al., 2007; Packard et al., 2016; Panaliappan et al., 2018; Tucker et al., 2010), Six1 (Chen et al., 2009; Ikeda et al., 2007), Meis1 (Toresson et al., 2000; Tucker et al., 2010), Pax7 (Murdoch et al., 2010; Stoykova and Gruss, 1994), and Ascl1 (Cau et al., 2002; Cau et al., 1997; Gordon et al., 1995; Guillemot et al., 1993). Despite detailed analyses based mostly upon loss-of-function mutations, there is still little insight into proliferative and lineage mechanisms that establish additional diversity within precursor classes defined by single molecular determinants. Precursors may acquire additional identities or lineage capacity based upon time of origin, transcriptional history, position in the OE, or local signals. Regulation of monoallelic, regionally-restricted expression of odorant receptor genes (reviewed by Monahan and Lomvardas, 2015) might rely upon this sort of additional precursor diversity, perhaps established at the outset of OE differentiation when position and signaling in the olfactory placode and frontonasal mass, a fairly circumscribed embryonic domain, can impart significant instruction for subsequent development (Balmer and LaMantia, 2005; LaMantia, 2020). OE precursor diversity established at early stages might endure to support mosaic molecular distinctions in the mature OE. Nevertheless, it remains unknown whether molecularly distinct precursors are initially subdivided into temporally distinct cohorts with specific proliferative and cell fate characteristics either during development or in the adult.

Progenitors identified by key transcription factors can be classified based upon responses to local signals or proliferative characteristics (Rawson and LaMantia, 2007; Tucker et al., 2010; Whitesides et al., 1998; Yang et al., 2018). It is unclear, however, whether such OE precursors are further subdivided based upon temporal distinctions, position, molecular identity, and lineage. We found a temporally distinct subset of early-generated differentially proliferative Ascl1+ precursors, likely derived from Pax7+ progenitors, restricted to a dorsomedial OE domain. These cells subsequently generate spatially segregated ORNs, as well as subsets of sustentacular cells and olfactory ensheathing glia. The progeny of these precursors may endure as a distinct lineage, or they may represent a transient cohort that gives rise to an initial, ephemeral subset of ORNs and supporting cells as the OE expands during fetal development.

## MATERIALS AND METHODS

### Animals

All animal procedures were reviewed and approved by the George Washington University Institutional Animal Care and Use Committee. *Sox2*^eGFP^ (Ellis et al., 2004), *Pax7*^Cre^(Keller et al., 2004), *Ascl1*^Cre-E^ (Battiste et al., 2007), and *Ai9:tdTomato*^fl^ reporter (Madisen et al., 2010) alleles were maintained on a C57Bl6N background. *Sox2* reporter and Cre driver alleles were transmitted paternally and *Ai9*^fl^reporter alleles maternally. Pregnancies were timed by date of vaginal plug detection after overnight matings as E0.5. Single injections of tamoxifen (10mg/ml; 100µl/dam) were given to pregnant dams carrying *Ascl1*^Cre-E+/-^:*Ai9* reporter^fl/-^litters. One subset of these dams received an injection of BrdU (50mg/kg body weight) on E11.5, 2 hours or 5 days prior to embryo collection; another subset received BrdU in their drinking water for 48 hours starting at E11.5, ending on E13.5, prior to collecting fetuses as E16.5. Dams were sacrificed by cervical dislocation. E11.5 and E16.5 embryos/fetuses were fixed by immersion in 4% paraformaldehyde overnight at 4°C, and fetal membranes collected for genotyping.

### Tissue processing and immunohistochemical analysis

Fixed embryos were rinsed in PBS, pH 7.4, transferred sequentially to 10%, 20%, and 30% sucrose in 0.1M phosphate buffer (PB) for cryoprotection, embedded in 2.5% agar in 30% PB-sucrose to position them consistently, and then flash frozen in cryoembedding media (OCT) using liquid nitrogen-cooled 2-methyl butane. Blocks were sectioned at 10 µm, serial sections mounted onto glass slides that were stored at -20°C before immunostaining. Primary antibodies against tdTomato/red fluorescent protein (RFP; Abcam, rabbit), Pax7 (Developmental Hybridoma Studies Bank, mouse), Meis1 (Abcam, rabbit), Ascl1 (BD Bio, mouse; Chemicon, rabbit), Six1 (Proteintech, rabbit; Atlas, mouse), eGFP (Abcam), pSMAD (Millipore, mouse), BrdU (BD, mouse; Novus, rat), PH3 (Cell Signaling; rabbit), βIII-tubulin (Aves, chicken), NCAM (Millipore, rat), and OMP (Pierce Biotech; chicken) followed by 2° antibodies appropriate for single, double or triple label as described previously (Karpinski et al., 2016). Images were collected on a Leica Tiling epifluorescence or Zeiss 710 confocal microscope.

### Quantitative analysis of OE cell classes *in vivo*

We quantified the distribution of OE progenitors and neurons labeled with multiple markers in the E11.5 OE, divided into 10 sectors, each representing 1/10 of the inner OE perimeter (Tucker et al, 2010; **Figure 1H**_**1-3**_). To normalize analyses of individual animals from multiple litters (4 to 6 embryos analyzed/marker), DAPI-labelled cells were also counted in each sector (n marker+ cells/n DAPI+ cells/sector), and results for each molecular marker were calculated as a proportion, rather than absolute number, of cells in each sector.

**Figure 1:**
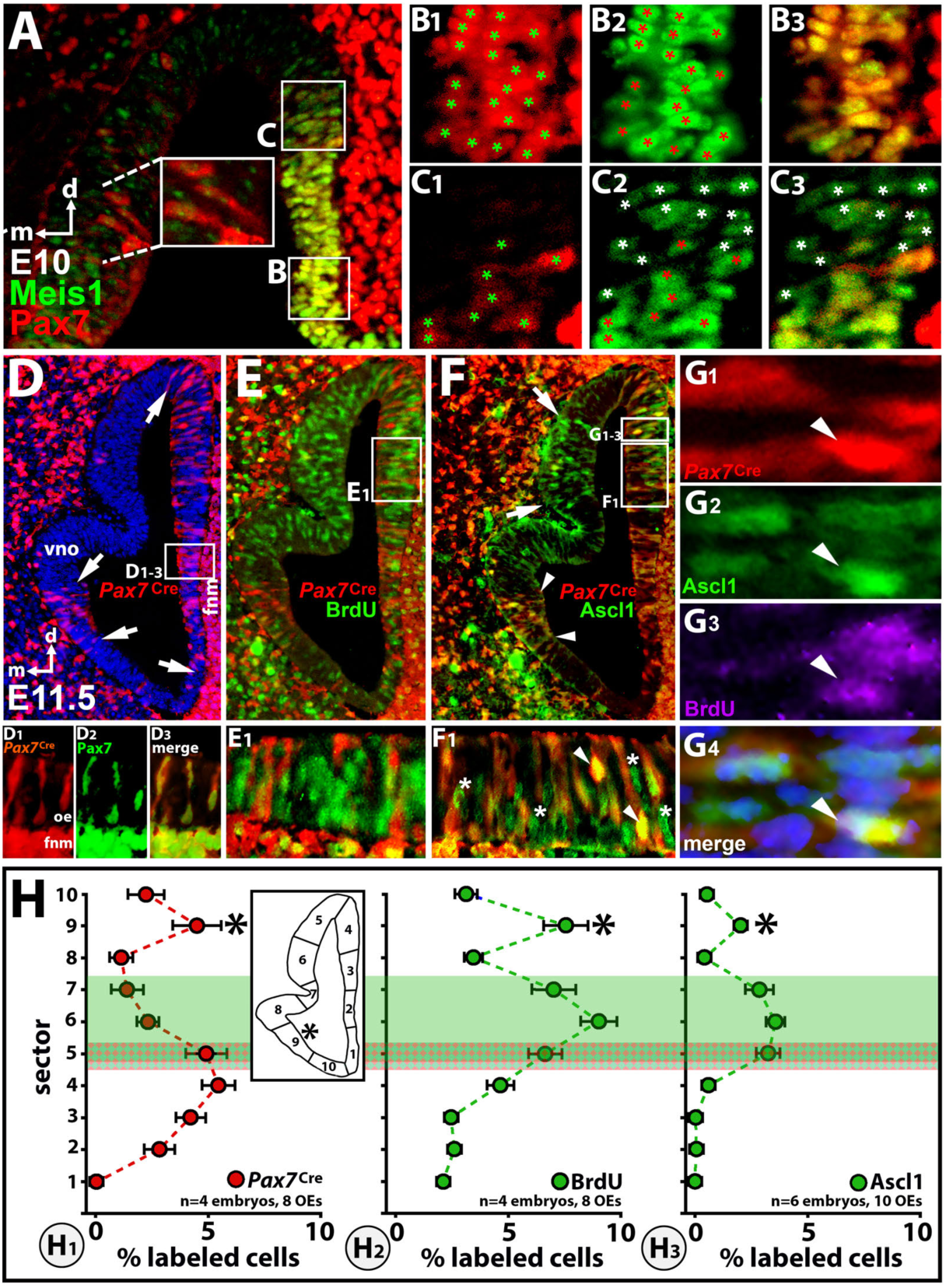
Distribution and identities of early Pax7+ olfactory epithelial (OE) progenitors. **A**) Pax7+ cells (immunolabeled; red), coincide with Meis1+ cells (immunolabeled; green) in the E10.0 lateral OE (coronal section; d=dorsal, m=medial). A small population of Pax7+/Meis1-cells is also seen in the ventromedial OE (dotted lines, box). **B1-3**) Nearly complete registration of Pax7+ (red label, green asterisks; *panel* **1**) and Meis1+ (green label, red asterisks; *panel* **2**) cells in the ventrolateral OE (merged; *panel* **3**). **C1-3**) Only a subset of dorsolateral Pax7+ cells (*panel* **1**) are co-labeled for Meis1+ (*panel* **2**). The single-labeled Meis1+ cells are indicated by white asterisks (merged; *panel* **3**). **D**) Cells labeled by a *Pax7*^Cre^-recombined floxed tdTomato reporter (*Pax7*^Cre^+) in the E11.5 OE (coronal section). *Pax7*^Cre^+ cells are most frequent in the lateral OE (arrows, right), and frontonasal mesenchyme (fnm). There is also a small domain of *Pax7*^Cre^+ cells in the medial OE, ventral to the presumptive vomeronasal organ (vno; arrows). **D1-3**) Registration of *Pax7*^Cre^+ (red) and Pax7+ (green) cells in E11.5 lateral OE and fnm. **E**) Relationship of *Pax7*^Cre^+ cells to proliferating OE precursors labeled acutely with BrdU, which are most frequent in the dorsal OE. **E1**) Lateral OE *Pax7*^Cre^+ cells are distinct from BrdU+ cells. **F**) Lateral *Pax7*^Cre^+ cells are segregated from dorsomedial Ascl1+ cells (arrows); however, *Pax7*^Cre^+/Ascl1+ cells are seen in the dorsolateral OE (box, **G1-4**) and a distinct ventromedial OE zone (arrowheads). **F1**) In the dorsolateral OE, a subset of *Pax7*^Cre^+ cells are also Ascl1+ (arrowheads); however, multiple Ascl1+ cells in this region are *Pax7*^Cre^-(asterisks). **G1-4**) In the dorsolateral OE there are occasional *Pax7*^Cre^+ cells (asterisk, **G1**) are also Ascl1+ (asterisk, **G2**) and BrdU+ (asterisk, **G3;** merged, **G4). H**) Relative frequency of *Pax7*^Cre^+ (**H1**), BrdU+ (**H2**) and Ascl1+ cells (**H3**), all as a proportion of total DAPI+ cells, measured in 10 linearly equivalent sectors, based upon the inner perimeter length of the OE (*inset*, left). The numbers of embryos (‘n”) and OE’s counted for each marker are provided in each of the graphs; errors bars reflect standard error of the mean for each point. The red shading indicates sectors where *Pax7*^Cre^+ cells are more frequent; green shading indicates sectors where BrdU+ and Ascl1+ cells are more frequent. There is a small zone of overlap (darker shading; at sector 5) The asterisk indicates a second peak (sector 9) where *Pax7*^Cre^+, BrdU+ and Ascl1+ cell frequency is elevated, in register with the ventromedial domain (arrows and arrowheads in panels **D** and **E**).

### Pair Cell Assays

*Sox2*^eGFP^ reporter expression was confirmed via fluorescence microscopy in E10.5 embryos and frontonasal processes were microdissected (n=4 independent experiments/4 litters). The complete OE was isolated, and then divided into medial and lateral samples. These samples were dissociated as described previously (Lehtinen et al., 2011; Shen et al., 2002; Tucker et al., 2010). Dissociated cells were plated at low density (35 cells/µl; 14 µl total volume/well) on poly-D-lysine coated Terasaki plates. Equivalent numbers of medial and lateral cells were plated initially, and similar numbers of wells were analyzed for each side subsequently. Cultures were incubated for 21 hours at 37°C with 5% CO_2_, fixed and immunolabeled for Sox2 (progenitor marker) and βIII-tubulin (neuronal marker), as well as DAPI to identify nuclei. Pairs of cells in individual wells were identified based upon DAPI labeling as well as eGFP fluorescence based upon apposition of two cells, isolated from any other cells. For each isolated DAPI-labelled pair, expression of eGFP, progenitor, and neural markers was visualized and scored.

## RESULTS

### A distinct population of Pax7+/Meis1+ lateral OE progenitors

We previously defined a population of Meis1+/Pbx1+/low-Sox2+ slowly dividing precursors in the E11.5 lateral OE (l-OE; Tucker et al., 2010). Pax7+ precursors in the nascent OE, defined both immunocytochemically and via *Pax7*^Cre^-mediated recombination (Murdoch et al., 2010) can also generate multiple OE cell types. We asked whether Meis1+ and Pax7+ cells define a single, early l-OE precursor population. Meis1+ and Pax7+ cells are found primarily in the l-OE at E10.5 (**Figure 1A**); however, there are also Pax7+/Meis1-cells in the medial/ventral OE (**Figure 1A**, box). l-OE Pax7+ and Meis1+ cells overlap nearly completely (**Figure 1B**_**1-3**_, asterisks); however, there is a small subset of Meis1+/Pax7-dorsal l-OE cells (**Figure 1C**_**1-3**,_ asterisks). Most *Pax7*^Cre^+ cells (**Figure D**_**1**_) can also be labeled for Pax7+ (**Figure D**_**2**,**3**_). This suggests very little lineage progression by Pax7+ cells by E11.5. Consistent with limited lineage progression for Pax7+/Meis1+ and *Pax7*^Cre^+ cells, acutely labeled BrdU+ cell frequency is lowest in l-OE regions where that of *Pax7*^Cre^+ cells is highest **(Figure 1E**,**H**_**1**,**2**_);there are few if any *Pax7*^Cre^+/BrdU+ cells (**Figure 1E**_**1**_), and PH3+ cell frequency is also lower in the l-OE (see **Figure 2**). A small population of *Pax7*^Cre^+ cells, however, coincide in the dorsal l-OE with Ascl1+ cells (**Figure 1F, F**_**1**_). Ascl1+ cells are thought to be a key class of more rapidly dividing OE precursor (Gordon et al, 1995; Cau et al, 1997; 2000; Tucker et al, 2010). Infrequent Ascl1+/ *Pax7*^Cre^+/ BrdU+ cells are found in this dorsal l-OE region (box, **Figure 1F**; **Figure 1G**_**1-4**_, asterisks). We quantified these impressions of distinct precursor distribution using a geometric sector analysis we described previously that permits assessing regional cell frequencies independent of size and shape variations of individual developing OEs (Tucker et al., 2010). In addition, marker-labeled cell frequency is calculated as a proportion of all cells in each sector, based upon counts of DAPI+ cells. *Pax7*^Cre^+ cell frequency is highest in l-OE (sectors 3,4; **Figure 1H**_**1**_). BrdU+ and Ascl1+ cell frequency is maximal in dorsomedial sectors (sectors 6,7; **Figure 1H**_**2**,**3**_).

**Figure 2:**
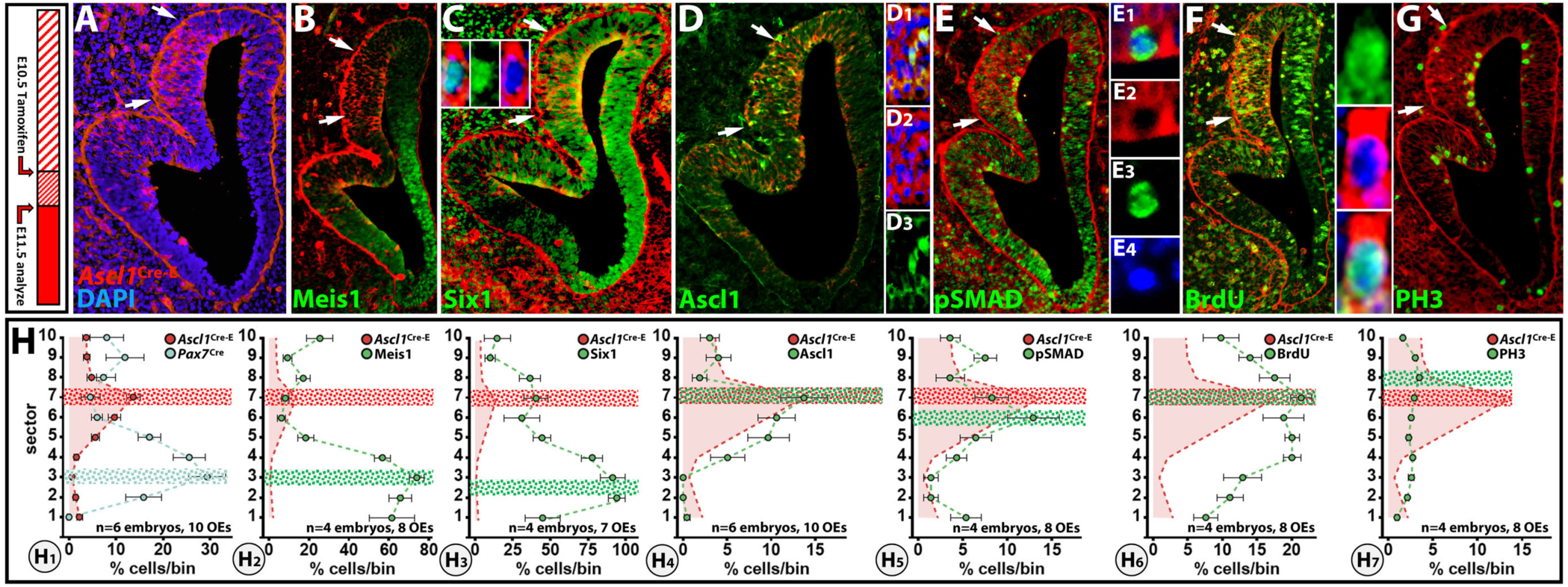
A distinct zone of highly proliferative Ascl1+ descended OE precursors. ***Far left***: the schedule of tamoxifen injection and embryo collection used to generate the data. **A**) After initiating recombination with a single E10.5 tamoxifen injection, there is a population of *Ascl1*^Cre-E^+ cells (arrows) concentrated in the dorsomedial OE. In subsequent graphs, a “ghost” image of the Ascl1Cre-E+ cell distribution, scaled to the cell frequency of the additional cell population, is shown by the red hatched curve. **B**) This population is completely segregated from Meis1+ lateral OE progenitors. **C**) Six1+ cells are seen throughout the lateral and dorsomedial OE, including in the region where *Ascl1*^Cre-E^+ cells are found. Some *Ascl1*^Cre-E^+ cells are also Six1+ (*inset*). **D**) A small population of Ascl1+ (protein) cells is found within the dorsomedial OE region where *Ascl1*^Cre-E^+ cells are concentrated. **D1-3**) Ascl1+ cells are a subset of the dorsomedial *Ascl1*^Cre-E^+ population. **E**) Phosphorylated SMAD 1/5/8 (pSMAD)-labeled nuclei are found primarily in apical *Ascl1*^Cre-E^+ dorsomedial OE cells. **E1-4**) An apical *Ascl1*^Cre-E^+ dorsomedial OE cell with a pSMAD+ nucleus. **F**) Dorsomedial *Ascl1*^Cre-E^+ cells coincide with actively proliferating BrdU+ cells, and many *Ascl1*^Cre-E^+ cells are also BrdU+ (inset). **G**) PH3+, presumably actively mitotic, cells are more frequent in the dorsomedial domain where *Ascl1*^Cre-E^+ cells are found. **H**) Quantitative analysis of frequency of Meis1+, Six1+, Ascl1+, pSMAD+, BrdU+, and PH3+ single-labeled cells, *Ascl1*^Cre-E^+ cells, and double labeled cells throughout the OE using the sector analysis method (see **Figure 1H**). Hatched red and green bars indicate maximal frequency for the relevant cell classes.

There is an apparent transition zone where frequency of all 3 markers increases (hatched region, **Figure 1H**). Thus, by E11.5, there is a minimally proliferative population of Pax7+, predominantly Meis1+ l-OE precursors and a small number of actively proliferating Ascl1+/BrdU+ Pax7+ descended cells in the dorsal l-OE.

### Intermediate OE precursors and their neuronal progeny

Previous observations suggest that Ascl1+ cells divide rapidly and asymmetrically to generate substantial numbers of ORNs (Guillemot et al., 1993; Murray et al., 2003). The distribution of OE Ascl1+ cells, their coincidence with markers of proliferative activity, and their fates, however, are variable and dynamic (Cau et al., 2002; Gordon et al., 1995; Tucker et al., 2010). To more clearly establish the derivation, identity, and fate of the earliest OE Ascl1+ precursors we used conditional Cre-recombinase driven by the endogenous *Ascl1* promoter (*Ascl1*^Cre-ERT^) at E10.5 to identify progeny at E11.5. These *Ascl1*^Cre-ERT^-recombined cells (*Ascl1*^Cre-E^+) accumulate in the dorsal medial-OE at E11.5 (m-OE; **Figure 2A**, arrows) coincident with Ascl1+ as well as BrdU+ cells (see **Figure 1**). This region is distinct from l-OE regions where Pax7+, *Pax7*^Cre^+ cells (see **Figure 1**), Meis1+ (**Figure 2B**), and Six1+ cells (**Figure 2C**), which are thought to be associated with general cranial placode precursors (Horie et al., 2018; Schlosser et al., 2014), are found (see also **Figure 2H**_**1-3**_). E11.5 *Ascl1*^Cre-E^+ cells overlap extensively with Ascl1+ protein cells (**Figures 2D**; **Figure 2H**_**4**_); however, *Ascl1*^Cre-E^+ cells are apparently more frequent than Ascl1+ or Ascl1+/*Ascl1*^Cre-E^+ cells (**Figure 2D**_**1-3;**_**Figure 2H**_**4**_).

We next asked whether there was a relationship between BMP or other TGF-β signaling associated with modulation of Ascl1 expression or additional transcription factors (Gokoffski et al., 2011; Shou et al., 1999; Wu et al., 2003) based upon the distribution of pSMAD 1/5/8 (pSMAD+), key intermediates for Tgfβ signaling in neural progenitors (Dincer et al., 2013; Hegarty et al., 2013). pSMAD+ cells are most frequent where Ascl1+ and *Ascl1*^Cre-E^+ cells are found (**Figure 2E**; **Figure 2H**_**4**,**5**_). Thus, a mosaic of Ascl1+ and *Ascl1*^Cre-E^+ cells in the dorsal m-OE apparently responds to Tgfβ signals, perhaps to regulate Ascl1 levels (Shou et al., 1999) or OE precursor neurogenic status (Gokoffski et al., 2011). Proliferative (BrdU+) frequency is highest in dorsal l-OE as well as dorsal m-OE (**Figure 2F**,**H**_**6**_ see also **Figure 1**); however, only a small proportion of m-OE *Ascl1*^Cre-E^+ cells are also BrdU+ (**Figure 2F, 2H)**. PH3+ presumed mitotic (von Bohlen und Halbach, 2011) cell frequency also increases in the m-OE, overlapping the Ascl1+ /*Ascl1*^Cre-E^+ domain, consistent with greater proliferative activity in this OE domain. Thus, temporally restricted conditional *Ascl1*^Cre-E^ lineage analysis identifies a spatially constrained dorsal m-OE precursor cohort with distinct molecular profiles and proliferative states.

### *Ascl1*^Cre-E^+ cells have distinct differentiation capacities

We next asked whether the E11.5 *Ascl1*^Cre-E^+ cohort generates nascent ORNs recognized by expression of βIII-tubulin (Roskams et al., 1998) as well as those initiating axon outgrowth from the OE toward the developing forebrain, recognized by expression of NCAM (Miragall et al., 1989). βIII-tubulin+ apparent ORNs (based upon cytological characteristics including an apically directed single dendrite) are seen occasionally in the domain where *Ascl1*^Cre-E^+ cells are also found; however, their frequency is apparently lower in this domain than flanking regions (**Figure 3A**_**1-2**_). Some βIII-tubulin+ cells are *Ascl1*^Cre-E^+ (**Figure 3B**_**1-3**_), while adjacent *Ascl1*^Cre-E^+ cells are not. Finally, βIII-tubulin+ ORNs appear more frequent in OE regions beyond the *Ascl1*^Cre-E^+ domain (**Figure 3A**_**1-2**_, **C**). βIII-tubulin also is localized to a substantial subset of *Ascl1*^Cre-E^+ cells in the migratory mass that encapsulates the nascent olfactory nerve (ON) at E11.5 (**Figure 3D**_**1-3**_). The proximity of βIII-tubulin+ cells and axons in the migratory mass makes it difficult to resolve whether the *Ascl1*^Cre-E^+ cells express βIII-tubulin or whether βIII-tubulin+/ *Ascl1*^Cre-E^ + ON axons are coincident with *Ascl1*^Cre-E^+ migratory mass cells (**Figure 3D**_**1-3**_).

**Figure 3:**
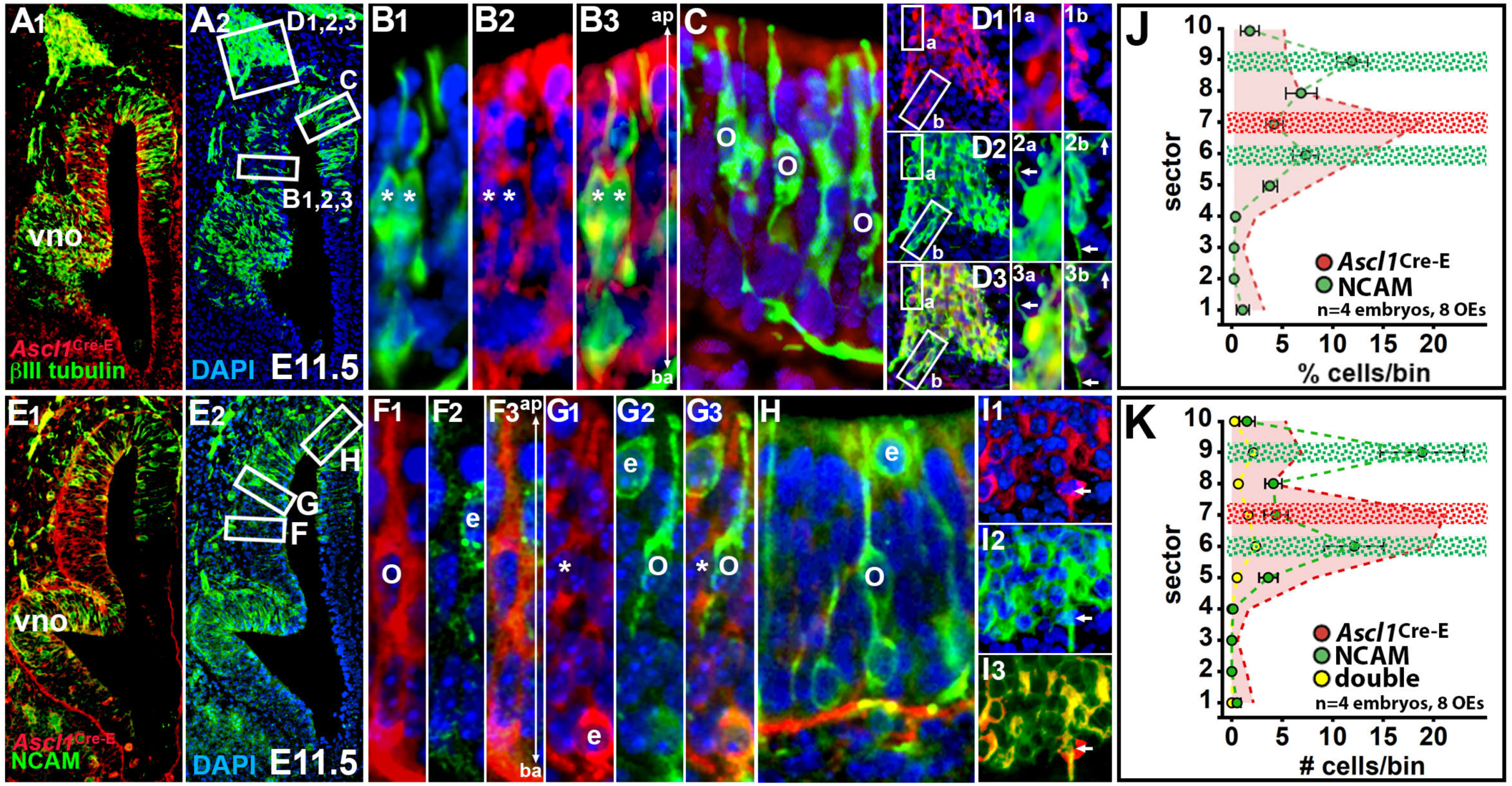
A distinct relationship between newly generated olfactory receptor neurons (ORNs) and *Ascl1*^Cre-E^+ precursors. **A1**,**2**) Early generated ORNs, labeled for the early neuronal marker bIII-tubulin, are distributed throughout the dorsolateral and dorsomedial OE. The frequency of bIII-tubulin+ presumptive ORNs seems somewhat attenuated in the region where *Ascl1*^Cre-E^+ cells are found. **B1-3**) Subsets of *Ascl1*^Cre-E^+ cells (asterisks) are bIII-tubulin+ ORNs with apically (ap) oriented dendrites. **C**) bIII-tubulin+ ORNs (o) in more lateral regions are not *Ascl1*^Cre-E^+ cells. **D1-3**) Ascl1Cre-E+ cells (1a,b), as well as apparent bIII-tubulin+ cells (2a,b) and axons constitute the migratory mass that constrains the nascent olfactory nerve as it extends through the frontonasal mesenchyme toward the ventral forebrain. **E**) A subset of NCAM+ ORNs are excluded from the dorsomedial region where *Ascl1*^Cre-E^+ cells are found. **F1-3**) *Ascl1*^Cre-E^ + cells with apically-oriented processes are seen in the dorsomedial OE; however, they are distinct from occasional NCAM+ cells within of *Ascl1*^Cre-E^+ domain. **G1-3**) At the margin of the dorsomedial domain where Ascl1Cre-E+ are concentrated, NCAM+ ORNs can be found adjacent to *Ascl1*^Cre-E^+ cells (asterisk). **H**) NCAM+ cells in the dorsolateral OE have essential characteristics of ORNs (o), including apically oriented apparent dendrites and apparent dendritic knobs (arrowhead), and can be found adjacent to Ascl1^Cre-E^+ epithelial cells (e). In addition, there are NCAM+ apical epithelial cells (e). **I1-3**) A substantial subset—but not all—of migratory mass cells in the frontonasal mesenchyme are *Ascl1*^Cre-E^+ (**I1**). NCAM+ cells and process (**I2**) coincide with the subset of Ascl1Cre-E+ migratory mass cells; but the overlap is not complete (**I3**). **J**) Sector plot of NCAM+ cell proportional frequency (to all DAPI+ cells/sector), with the distribution of *Ascl1*^Cre-E^+ cell proportional frequency provided (red curve) for comparison. **K**) Sector plot of absolute numbers of NCAM+ (green), *Ascl1*^Cre-E^+ (red) and NCAM+/ *Ascl1*^Cre-E^+ double labeled cells (yellow).

We next assessed the distribution of NCAM+ ORNs thought to be actively extending an axon (Miragall et al., 1989; Yamashita et al., 1998). NCAM+ ORNs are seen at frequencies similar to βIII-tubulin+ ORNs in the dorsal l-OE and ventral m-OE; however, they appear to be mostly excluded from the *Ascl1*^Cre-E^+ domain (**Figure 3E**, and **Figure 3J**, graph), and there are few NCAM+/ *Ascl1*^Cre-E^+ cells (**Figure 3F-H, K**). Some *Ascl1*^Cre-E^+ cells appear cytologically similar to nascent ORNs (**Figure 3F**_**1-3**_); however, there are also NCAM+ /*Ascl1*^Cre-E^+/-undifferentiated epithelial cells (**Figure 3G**_**1-3**_ and *inset*), adjacent to or occasionally within the region of *Ascl1*^Cre-E^+ precursors. In contrast, *Ascl1*^Cre-E^-/NCAM+ ORNs are fairly frequent in regions most distal to the *Ascl1*^Cre-E^+ precursor domain (**Figure 3H**).

Apparently, Ascl1+ cell-derived OE precursors remain segregated, and most do not immediately generate ORNs. Finally, *Ascl1*^Cre-E^+/Ascl1-cells and processes coincide with NCAM+ cells and processes in the migratory mass (**Figure 3I**_**1-3**_). *Ascl1*^Cre-E^ + cells are embedded in a broader domain of NCAM+ cells and processes (arrow, **Figure 3I**_**1-3**_). The registration of *Ascl1*^Cre-E^+ and NCAM+ cells and processes in the migratory mass is not as complete as for βIII-tubulin+ cells and process. This suggests that a subset of *Ascl1*^Cre-E^+ migratory mass cells are associated with actively growing ON axons, while others interact with additional ORN axons that may have reached their initial forebrain target.

### Distinct origins of frontonasal mesenchymal cells associated with the olfactory nerve

The apparent accumulation of *Ascl1*^Cre-E^+/Ascl1-cells in the migratory mass suggests that OE precursors contribute to the frontonasal mesenchyme (fnm), a mosaic of cells with distinct molecular identities, and signaling capacity (reviewed by LaMantia, 2020). Medial, and some dorsolateral, fnm cells express Six1 (**Figure 4A**), while lateral fnm cells express Pax7 (**Figure 4B**). Migratory mass cells, however, are distinct. They express neither Six1 nor Pax7; instead, they are *Ascl1*^Cre-E^+ (insets, **Figure 4A**,**B**). These *Ascl1*^Cre-E^+ migratory mass cells do not express Ascl1 protein (data not shown), which is limited primarily to m-OE cells (see **Figure 2**) nor do they express Pax7 protein (**Figure 4B**, *inset*). Thus, the initial cohort of migratory mass cells are apparently derived from early generated Ascl1+ OE precursors, consistent with previous observations that migratory mass cells originate primarily from the OE itself (LaMantia et al., 2000; Rawson et al., 2010). A small number of *Pax7*^Cre^+ medial fnm mesenchymal cells coincide with NCAM+ ORN axons or NCAM+ fnm cells (**Figure 4C**_**1**,**2**_). This suggests that *Ascl1*^Cre-E^+ migratory mass cells descend from Ascl1+ precursors derived from OE Pax7+ progenitors.

**Figure 4:**
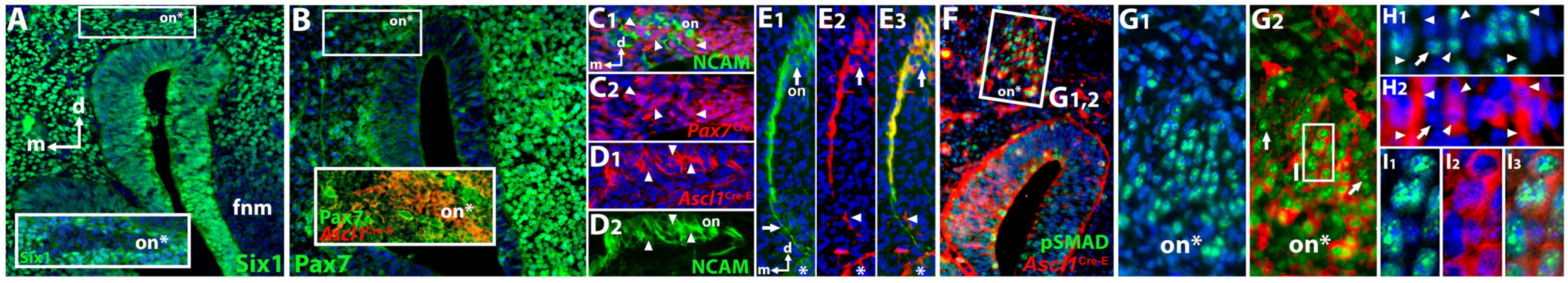
Distinctions in identity and lineage of olfactory nerve-associated frontonasal mesenchyme. **A**) Six1+ frontonasal mesenchymal (fnm) cells are concentrated in the medial and dorsolateral fnm, but attenuated in the lateral ventral fnm. In the dorsomedial fnm however, there are regions where Six1+cells are not seen (box and inset), presumably where the olfactory nerve (on*) is forming. **B**) Pax7+ (protein) cells are concentrated in the lateral fnm. In the mediodorsal fnm where fascicles of the olfactory nerve are presumably growing (on*) there are few, if any Pax7+ cells. In this mediodorsal region where Pax7+ cells are absent, *Ascl1*^Cre-E^+ cells define the migratory mass and associated olfactory nerve axon fascicles (box, *inset*, on*). **C1-2**) NCAM+ axons and cells of the nascent olfactory nerve (on; **C1**) and migratory mass are seen at the boundary of the medial and lateral fnm, coincident with *Pax7*^Cre^+ cells (arrowheads **C1, 2**). **D1-2**) In the mediodorsal fnm there are accumulations of *Ascl1*^Cre-E^+ cells and processes (arrowheads, **D1**) that predict where NCAM+ cells and processes that constitute the olfactory nerve (on) will be found. **E1-3**) Small fascicles of NCAM+ axons (**E1**; arrows) contact occasional *Ascl1*^Cre-E^+ cells (**E2**) as they exit the olfactory epithelium (asterisk). **F**) *Ascl1*^Cre-E^+ cells in the migratory mass as well as in the dorsomedial OE are also pSMAD+. These cells are segregated within the fnm in the region where the olfactory nerve (box, on*) coalesces as it extends toward the ventral forebrain. **G1-2**) pSMAD+ cells define a distinct, restricted subpopulation of fnm cells. pSMAD+ nuclei tend to be aligned in the region of the presumptive olfactory nerve (on*). Most pSMAD+ cells appear to be *Ascl1*^Cre-E^+ as well; however, there are some pSMAD+/*Ascl1*^Cre-E^-cells (arrows). **H1-2**) Arrowheads indicate regions where pSMAD+ fnm cells appear aligned, adjacent to *Ascl1*^Cre-E^+ axon fascicles. Some pSMAD+ cells do not appear to be *Ascl1*^Cre-E^+ (arrow). **I1-3**) Individual clusters of Ascl1^Cre-E^+ cells in the migratory mass appear to be pSMAD+ as well.

There is partial registration of *Ascl1*^Cre-E^+ cells and processes with those labeled for NCAM in the ON as it coalesces in the dorsomedial fnm (**Figure 4D**_**1-2**_, see also **Figure 3**). In addition, small fascicles of NCAM+ ORN axons that originate in the ventral m-OE coincide with a subset of *Ascl1*^Cre-E^+ cells in the medial fnm prior to coalescing with the migratory mass (**Figure 4E**_**1-3**_). Apparently, early generated *Ascl1*^Cre-E^+ cells establish and maintain a defined fnm territory that supports initial ORN axon growth to the forebrain (Miller et al., 2010; Whitesides and LaMantia, 1996). Finally, we asked whether *Ascl1*^Cre-E^+ cells in the migratory mass share characteristics with *Ascl1*^Cre-E^+ cells within the dorsomedial OE. There is a subset of pSMAD+/*Ascl1*^Cre-E^+ cells in the fnm associated with *Ascl1*^Cre-E^+ cells and processes (**Figure 4F**). The pSMAD labeling of these presumed migratory mass cells, parallels that of *Ascl1*^Cre-E^+ cells in the dorsomedial OE, and further indicates a role for Tgfβ/SMAD-mediated signaling in differentiation of the progeny of Ascl1+ OE precursors. Additional fnm cells are either only pSMAD+ or *Ascl1*^Cre-E^+ (**Figure 4G**_**1**,**2**_) and are found adjacent to apparent Ascl1Cre-E+ ORN axon fascicles (**Figure 4H**_**1**,**2**_). This suggests additional cellular diversity, beyond the early Ascl1+ lineage, within the migratory mass, perhaps including GnRH neurons migrating from the OE at this time (reviewed by Schwanzel-Fukuda et al., 1992; Wray, 2010). *Ascl1*^Cre-E^ labeling is seen in the peri-nuclear cytoplasm and proximal processes of a subset of pSMAD+ migratory mass cells (**Figure 4I**_**1-3**_). Apparently, cells descended from an initial cohort of Ascl1+ OE precursors, including those that are pSMAD+, contribute to the migratory mass. They delaminate from the dorsomedial OE and initiate or maintain Tgfβ responses as they support initial coalescence and growth of axons from nascent ORNs that are not derived from Ascl1+ precursors in the early developing OE.

### Identity and mode of division of distinct E11.5 OE progenitor lineages

The temporal, regional and lineage-associated diversity of early OE precursors suggests that these cells differ in their proliferative characteristics. We used a pair cell assay (Shen et al, 2002; Tucker et al, 2010; Lehtinen et al, 2011; Karpinski et al, 2021) to distinguish modes of division and fates of isolated E11.5 medial versus lateral OE precursors. A *Sox2*^eGFP^reporter expressed throughout the developing OE but not in the fnm (Ellis et al, 2004; Tucker et al, 2010) identifies presumed neural precursors from the dissected E11.5 medial and lateral OE (**Figure 5A**). Subsets of *Sox2*^eGFP^+ pairs express Six1 (**Figure 5B**), Pax7 (**Figure 5C**), Ascl1 (**Figure 5D**), and βIII-tubulin (**Figure 5E**). We analyzed labeled pairs in two ways: first, as a proportion of all pairs in which at least one cell is *Sox2*^eGFP^+ (minimally *Sox2*^eGFP^+). Medial OE precursors appear to be more proliferative based on recovery of more minimally *Sox2*^eGFP^+ medial versus lateral *Sox2*^eGFP^+ pairs from equivalent platings from 4 independent experiments (medial=1589 total *Sox2*^eGFP^ pairs; lateral=829 *Sox2*^eGFP^+ pairs). The proportions of medial versus lateral pairs labeled for at least one of the markers analyzed *in vivo* in this study, however, did not differ significantly (**Figure 5F**_**1**_). Next, we analyzed medial vs. lateral Six1+, Pax7+, Ascl1+, βIII-tubulin+ pairs as a proportion of minimally Sox2^eGFP^+ medial or lateral pairs. Six1+ medial pairs were more frequent (17% medial, 13% lateral; p≤0.04; Fisher Exact; **Figure 5F**_**2**_) and Pax7+ lateral pairs were more frequent (4% medial, 14% lateral; p≤0.00001; Fisher Exact; **Figure 5F**_**2**_). This reinforces the conclusion that medial and lateral precursors are distinct from one another in the context of molecular identities assessed in our *in vivo* analysis. In contrast, Ascl1+ and βIII-tubulin+ pairs, presumably “downstream” or neurogenic precursors, respectively, do not differ in frequency as a proportion of minimally Sox2eGFP+ pairs (**Figure 5F**_**2**_). We then analyzed Six1+, Pax7+, Ascl1+ or βIII-tubulin+ pairs as a proportion of all pairs that expressed at least one marker, thus excluding pairs with unknown molecular identity. Pax7+ lateral pairs are more frequent (13% medial, 38% lateral; p≤0.00001; Fisher Exact; **Figure 5F**_**3**_), as are Ascl1+ medial pairs (17% medial, 10% lateral; p≤0.03; Fisher Exact; **Figure 5F**_**3**_). This is consistent with enhanced proliferative Ascl1+ precursors in the medial OE, perhaps generated from a more slowly dividing Pax7+ lateral precursor population.

**Figure 5:**
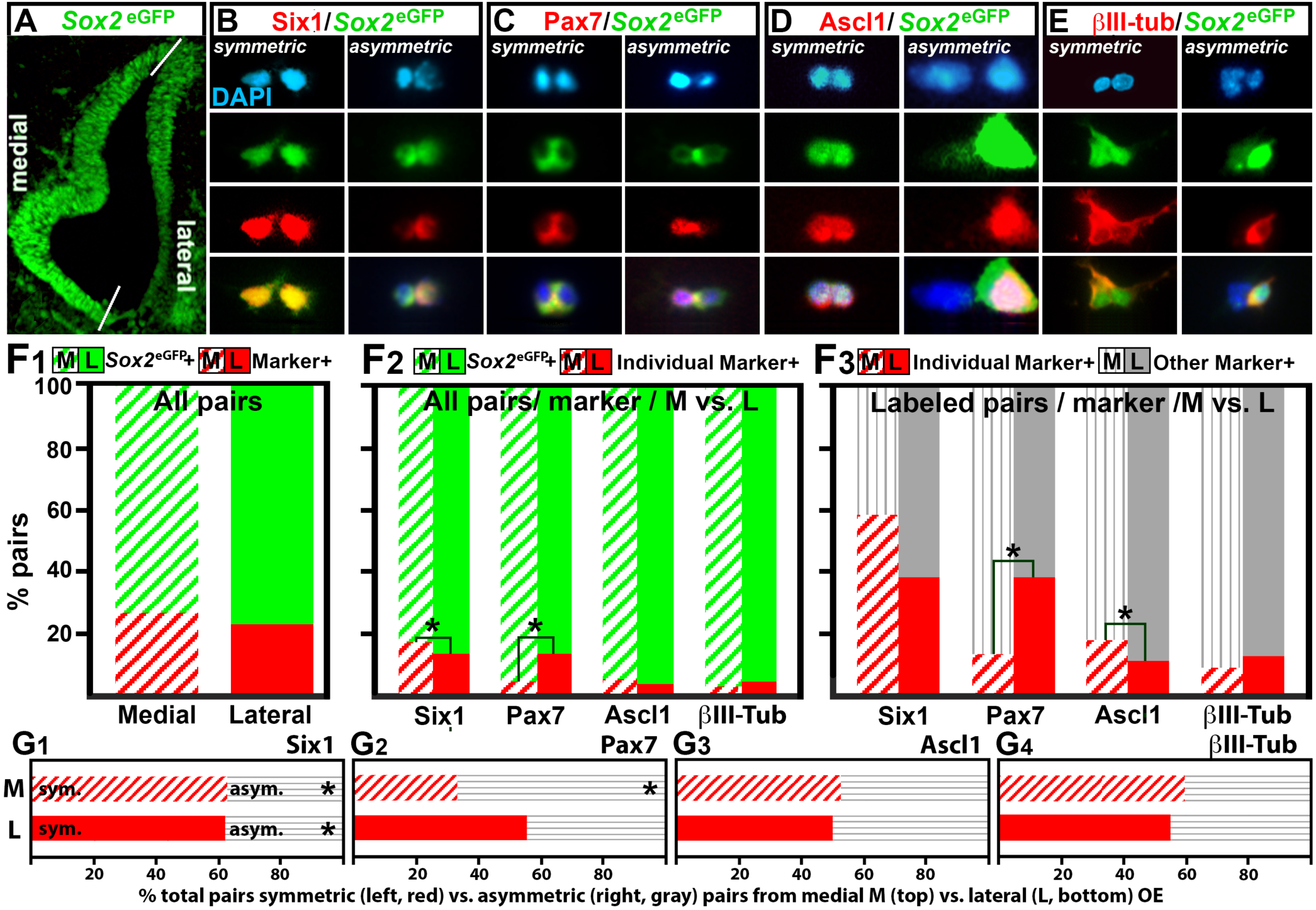
Molecular identity and OE position influence modes of precursor proliferation and division. **A**) A *Sox2*^eGFP^ reporter transgene labels lateral and medial OE cells in a graded fashion. Symmetric and asymmetric divisions of **B**) Six1+/*Sox2*^eGFP^+ precursor cell pairs recorded and scored in this analysis. **C**) Pax7+/*Sox2*^eGFP^+ cell pairs. **D**) Ascl1+/*Sox2*^eGFP^+ cell pairs. **E**) symmetric and asymmetric neurogenic divisions from Sox2eGFP+ progenitors identified with βIII-tubulin. **F**) Normalized frequency of medial versus lateral cell pairs. **F1**) The frequency of medial versus lateral pairs labeled by *Sox2*^eGFP^ and any of the four additional markers does not differ significantly. **F2**) The frequencies of Six1+ and Pax7+ pairs as a proportion of all *Sox2*^eGFP^+ medial versus lateral cell pairs differ significantly (asterisks, Six1+ M > Six1+ L, p ≤ 0.04; Fisher-Exact; Pax7+ M < Pax7+ L, p ≤ 0.001, Fisher-Exact). **F3**) Pax7+ pairs are significantly more frequent among lateral pairs (asterisk, p ≤ 0.0001; Fisher-Exact) and Ascl1+ pairs are more frequent among medial pairs (asterisk. p ≤ 0.03, Fisher-Exact) when frequency of medial versus lateral pairs is compared for each marker class as a proportion of all medial or lateral cells labeled for any of the four markers. **G**) Normalized frequency of symmetric versus asymmetric dividing cell pairs from the medial and lateral OE. **G1**) For medial and lateral Six1+ cell pairs, there are significantly more symmetric divisions (p ≤ 0.0001 medial; 0.03, lateral; Fisher Exact). **G2**) There are significantly fewer symmetric divisions among lateral Pax7+ pairs (p≤0.03, Fisher Exact); however, the frequency of symmetric versus asymmetric divisions for medial Pax7+ cell pairs do not differ significantly. **G3**) There is no significant difference in symmetric versus asymmetric divisions for medial or lateral Ascl1+ cell pairs. **G4**) There is no significant difference in neurogenic symmetric versus asymmetric divisions (βIII-tubulin+) for medial or lateral pairs.

Biases toward symmetric versus asymmetric division also distinguish subclasses of OE precursors, despite an overall equivalence of medial versus lateral symmetric and asymmetric divisions across minimally *Sox2*^eGFP^+ pairs. Six1+ medial as well as Six1+ lateral precursors divide symmetrically significantly more frequently (p≤0.0001, medial; 0.03, lateral; Fisher Exact; **Figure 5G**_**1**_). In contrast, Pax7+ medial precursors divide asymmetrically significantly more frequently (p≤0.03; Fisher Exact; **Figure 5G**_**2**_). Medial and lateral Ascl1+ precursors divide symmetrically or asymmetrically with approximately equivalent frequency (**Figure 5G**_**3**_), consistent with a mixture of Ascl1+ self-renewing and neurogenic transit-amplifying precursors, regardless of location. The neurogenic yield (βIII-tubulin+) from *Sox2*^eGFP^+ medial and lateral asymmetric versus symmetric divisions is equivalent (**Figure 5G**_**4**_).

Thus, modes of division of subsets of OE precursors reflect both initial OE position and molecular identity. These differences, in the context of diversity introduced by temporal distinctions among molecularly defined lineages modulated by local signals, may influence the capacity of subsets of OE precursors to generate specific OE differentiated cell classes.

### Differentiated cell fates of a distinct precursor class in the developing OE

We next asked whether the E10.5-recombined (E10.5-R) *Ascl1*^Cre-E^ + temporal/lineage cohort gives rise to identifiable, spatially-restricted differentiated OE cell classes at E16.5, as the OE grows and acquires a mature turbinate morphology (**Figure 6A**_**1-4**_). At E16.5, E10.5-R *Ascl1*^Cre-E^+ cells include a subpopulation of ORNs, sustentacular cells, and ensheathing glia in the anterior OE. From the most anterior (**Figure 6A**_**1**,**2**_) to mid-anterior levels of the E16.5 OE (**Figure 6A**_**3**,**4**_), E10.5-R *Ascl1*^Cre-E+^ cells are distributed discontinuously (**Figure 6A**_**1**_, *inset*). E10.5-R *Ascl1*^Cre-E^+ cells are most prominent in a medial/ventral domain in the mid-anterior E16.5 OE (**Figure 6A**_**2**,**3**_). Posterior to this region, a patchy distribution returns in medial turbinates (**Figure 6A**_**4**_). The cytology of E10.5-R *Ascl1*^Cre-E^+-derived cells varies in the E16.5 OE, as does basal/apical location (**Figure 6B, C**), and there are few, if any E10.5-R *Ascl1*^Cre-E^+ cells in the respiratory epithelium (**Figure 6D**). E10.5-R *Ascl1*^Cre-E^+ OE cells at E16.5 appear to be ORNs (o), sustentacular cells (s), and far less frequently, precursors (p). In addition, in the lamina propria adjacent to the OE, there are individual E10.5-R *Ascl1*^Cre-E^+ cells as well as *Ascl1*^Cre-E^+ axon fascicles (**Figure 6C**, asterisks; **Figure 6E**, arrowheads). To assess the retention of proliferative capacity versus neurogenesis of the E10.5-R *Ascl1*^Cre-E^+ cohort, we gave E10.5-R dams BrdU in their drinking water for 48 hours starting at E11.5 and assessed heavily labeled OE cells in their E16.5 fetuses. If a substantial number of E10.5-R *Ascl1*^Cre-E^+ cells are transit amplifying precursors that generate substantial numbers of post-mitotic ORNs via multiple divisions, few E16.5 ORNs would be terminally labeled due to serial dilution of the label in successive proliferative daughter cells. This is indeed the case; there are few *Ascl1*^Cre-E^+ORNs in the E16.5 OE that are heavily labeled by BrdU (**Figure 6F**). Some *Ascl1*^Cre-E^+/BrdU+ cells in the mid-basal or basal OE (**Figure 6G**,**H**) may also be sustentacular supporting cells or “label-retaining” precursors (Morshead et al., 1994; Tucker et al., 2010). Thus, the E10.5-R *Ascl1*^Cre-E^+ lineage cohort generates multiple OE cell classes in distinct proliferative states by E16.5.

**Figure 6:**
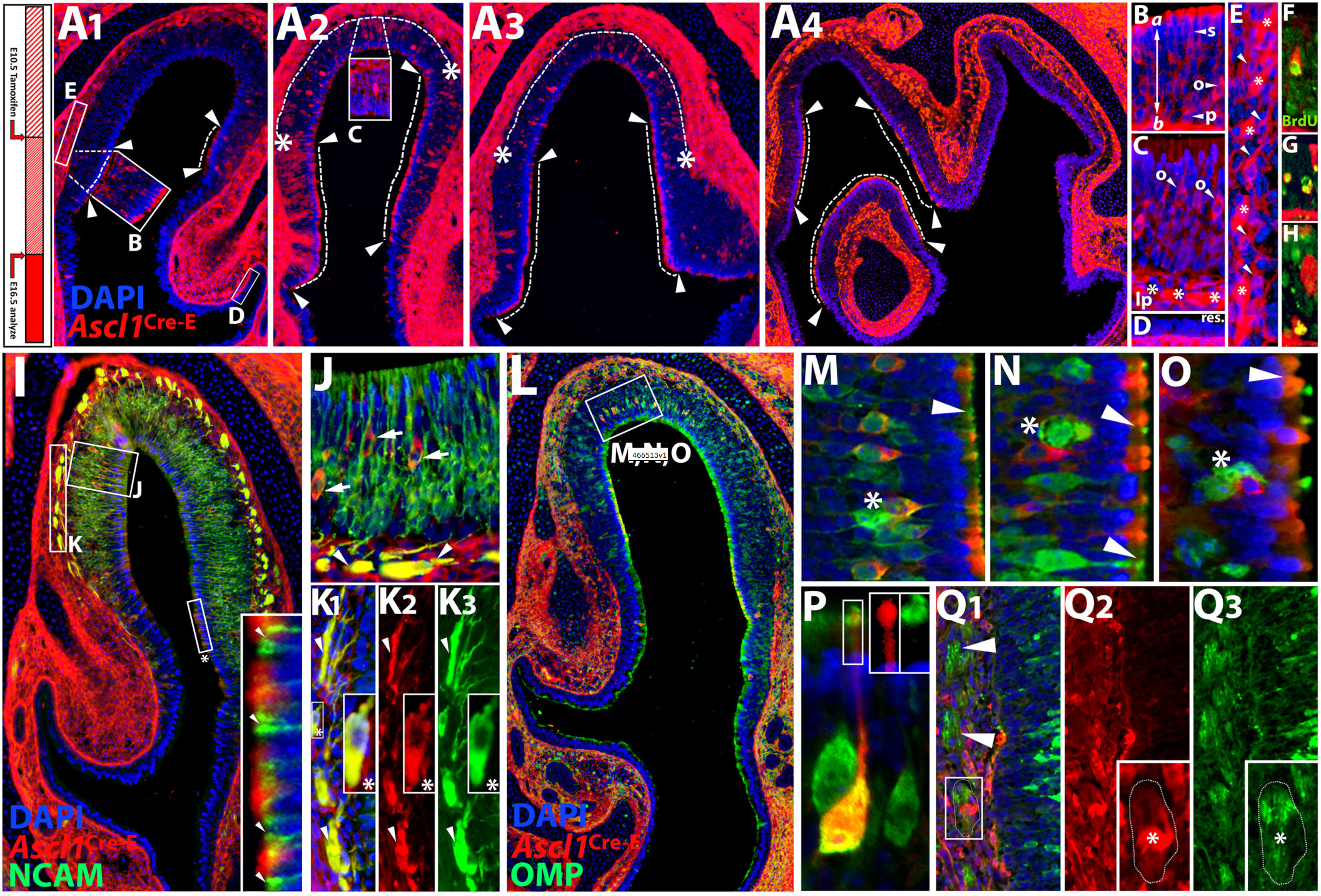
Restricted distribution and diverse fates of E10.5 recombined (E10.5-R) *Ascl1*^Cre-E^+ precursor cohort in the OE at E16.5. **A1-4**) Series of sections showing the distribution of E10.5-R *Ascl1*^Cre-E^+ cells. The dotted lines between arrowheads indicate regions where apparent sustentacular/supporting cells and their processes predominate, primarily in the more ventral aspect of the anterior OE at the transition from respiratory epithelium. The dotted lines between asterisks indicate regions where apparent E10.5-R *Ascl1*^Cre-E^+ ORNs predominate, primarily in dorsal anterior OE regions. The areas boxed in A1-4 are enlarged in **B-E. B**) A region where E10.5-R *Ascl1*^Cre-E^+ sustentacular/supporting cells (s, arrowhead) are most frequent. Many *Ascl1*^Cre-E^+ sustentacular cells have apical extensions that end in club-like processes at the apical (*a*)/lumenal surface. Other E10.5-R *Ascl1*^Cre-E^+ cells at more basal positions (*b*) may be precursor cells (p, arrowhead) or nascent ORNs (o, arrowhead). **C**) A region where apparent *Ascl1*^Cre-E^+ ORNs (arrowheads) that extend dendrites toward the apical/lumenal surface predominate. In the lamina propria (lp) reporter labeling (asterisks) likely reflects ORN axon fascicles and possibly ensheathing cells that coalesce after the axons exit the OE. **D**) There are no E10.5-R *Ascl1*^Cre-E^+ cells in the respiratory epithelium (res). **E**) In the lamina propria distal to the OE, E10.5-R *Ascl1*^Cre-E^+ cells and processes accumulate. E10.5-R *Ascl1*^Cre-E^+ cells (arrowheads) appear to be olfactory ensheathing cells that envelop fascicles of ORN axons as they grow toward the olfactory nerve. **F-H**) Occasional E11.5-13.5 chronic BrdU+/ E10.5-R *Ascl1*^Cre-E^+ cells are seen at mid to basal levels in the E16.5 OE. **F**) E10.5-R *Ascl1*^Cre-E^+ apparent ORN. **G**) BrdU+/ E10.5-R *Ascl1*^Cre-E^+ cells also have sustentacular/epithelial morphology. **H**) An apparent BrdU+**/**E10.5-R *Ascl1*^Cre-E^+ precursor in the basal OE. **I**) Infrequent E10.5-R *Ascl1*^Cre-E^+ ORNs among the far more numerous NCAM+ ORNs. **J**) Most, if not all, E10.5-R *Ascl1*^Cre-E^+ ORNs are NCAM-. In the lamina propria, E10.5-R *Ascl1*^Cre-E^+ cells and processes coincide with NCAM+ ON axon fascicles (arrowheads). **K1-3**) Small fascicles of NCAM+ axons exit the OE, extend through the lamina propria and join larger accumulations of E10.5-R *Ascl1*^Cre-E^+ cells and processes, which are also NCAM+. Apparent E10.5-R *Ascl1*^Cre-E^+ ensheathing cells (asterisk) are NCAM+ as well. E10.5-R *Ascl1*^Cre-E^+/NCAM+ processes accumulate where no cell nuclei are seen (arrowheads). **L**) E10.5-R *Ascl1*^Cre-E^+ ORNs are found among OMP+ ORNs in the E16.5 dorsal anterior OE. **M-O**) Some E10.5-R *Ascl1*^Cre-E^+ ORNs are OMP+; however, some are not. E10.5-R *Ascl1*^Cre-E^+/OMP+ ORNs can be immediately adjacent to those that are OMP+ only (asterisk, **M**). In addition, some E10.5-R *Ascl1*^Cre-E^+ only ORNs are immediately adjacent to OMP+ only ORNs (asterisks, **N, O**). The dendrites and knobs of OMP+ ORNS are often intercalated between E10.5-R *Ascl1*^Cre-E^+ club-like endings of presumed sustentacular cells (arrowheads, **M**,**N**,**O**). **P**) Occasional differentiated E10.5-R *Ascl1*^Cre-E^+/OMP ORNs with apical dendrites and dendritic knobs (*insets*) in the dorsal anterior E16.5 OE,are often adjacent to OMP+ only ORNs. **Q1-3**) Subsets of OMP+ ORN axon fascicles coalesce with apparent E10.5-R *Ascl1*^Cre-E^+ olfactory ensheathing cells and processes in the lamina propria (arrowhead, and *insets*, **Q2**,**3**).

To assess the relationship between E16.5 ORNs and the E10.5-R *Ascl1*^Cre-E^+ cohort, we labeled differentiating E16.5 E10.5-R *Ascl1*^Cre-E^+ for NCAM as well as OMP to distinguish likely actively differentiating ORNs with axons growing toward the olfactory bulb (OB) versus ORNs whose axons have reached the OB (Mont-Graziadei et al, 1980). NCAM+ ORN apical dendrites are intercalated among subsets of E10.5-R *Ascl1*^Cre-E^+ sustentacular cells (**Figure 6I, *inset***). Occasional E10.5 R *Ascl1*^Cre-E^+/NCAM-ORNs are found among a much larger population of NCAM+ ORNs (**Figure 6J**). There is extensive, but not complete registration of NCAM+ axon fascicles in the lamina propria with *Ascl1*^Cre-E^+ cells and processes (**Figure 6J**, arrowheads; **Figure 6K**_**1-3**_). Apparently, some E10.5-R *Ascl1*^Cre-E^+ lamina propria cells are OE ensheathing cells that interact with E10.5-R *Ascl1*^Cre-E^+/NCAM+ ORN axons. *Ascl1*^Cre-E^+/OMP+ ORNs are a subset of OMP+ ORNs (**Figure 6L-N**). Additional ORNs may be generated from Pax7+/Meis1+ progenitors (see **Figures 1, 3**) or a later-defined temporal/spatial cohort of Ascl1+ precursors. Subsets of OMP+/*Ascl1*^Cre-E^-distal dendrites are segregated from E10.5-R *Ascl1*^Cre-E^+ sustentacular cell apical processes (**Figure 6O**, arrowheads). Nevertheless, there are E10.5-R *Ascl1*^Cre-E^+/OMP+ ORNs with apical dendrites and nascent dendritic knobs, often adjacent to OMP+/*Ascl1*^Cre-E^-ORNs (**Figure 6O**, asterisks; **Figure 6P**, *insets*). Finally, we assessed the relationship between E10.5-R *Ascl1*^Cre-E^+ cells and processes in the lamina propria and OMP+ axons (**Figure 6Q**_**1-3**_). There is less overlap of OMP+ fascicles and *Ascl1*^Cre-E^+ versus NCAM+ cells and processes (**Figure 6Q**_**1**_). Instead, *Ascl1*^Cre-E^+ cells are often adjacent to OMP+ axon fascicles (**Figure 6Q**_**2**,**3**_, *insets*). Thus, the E10.5-R *Ascl1*^Cre-E^+ cohort generates spatially segregated subsets of ORNs with a broad range of differentiated states, as well as sustentacular cells, occasional precursors, and ensheathing cells that associate with subsets of ORN axons as they coalesce in the nascent ON.

### Ascl1+ OE progenitors and olfactory ensheathing cells

At E11.5, E10.5-R *Ascl1*^Cre-E^+ cells constitute a substantial subpopulation of migratory mass cells that interact with ORN axons as they extend toward the ventral forebrain in the fnm. By E16.5, E10.5-R Ascl1Cre-E+ ensheathing cells persist, closely associated with subsets of primarily NCAM+ axons. To further define the relationship E10.5-R *Ascl1*^Cre-E^+ fnm progeny and ORN axons, we analyzed at higher resolution the distribution of NCAM+ and OMP+ ORN axon fascicles with E10.5-R *Ascl1*^Cre-E^+ cells at E16.5 (**Figure 7A-D**). NCAM+ axon fascicles coincide with chains and aggregates of *Ascl1*^Cre-E^+ cells in the lamina propria as they extend toward larger fascicles that will define the ON (**Figure 7A**). In cross section, apparent processes of *Ascl1*^Cre-E^+ cells define septa that divide sub-fascicles of NCAM+ ORN axons (**Figure 7B, C**). When individual axon fascicles originating from NCAM+/*Ascl1*^Cre-E^-ORNs are visualized as they enter the lamina propria, tightly fasiculated NCAM+ ORN axons and *Ascl1*^Cre-E^+ cells and processes are often distinct (**Figure 7D**_**1-5**_; brackets vs. arrowheads). While some NCAM+ axon fascicles are apparently invested by *Ascl1*^Cre-E^+ cells and processes, other NCAM+ fascicles and *Ascl1*^Cre-E^+ cells/processes are not in complete registration (compare brackets and arrowheads, **Figure 7D**_**2**,**3**_). *Ascl1*^Cre-E^+ cells align with and envelop NCAM+ fascicles; however, the processes of the *Ascl1*^Cre-E^+ cells extend beyond the axon bundles (**Figure 7D**_**4**,**5**_).

**Figure 7:**
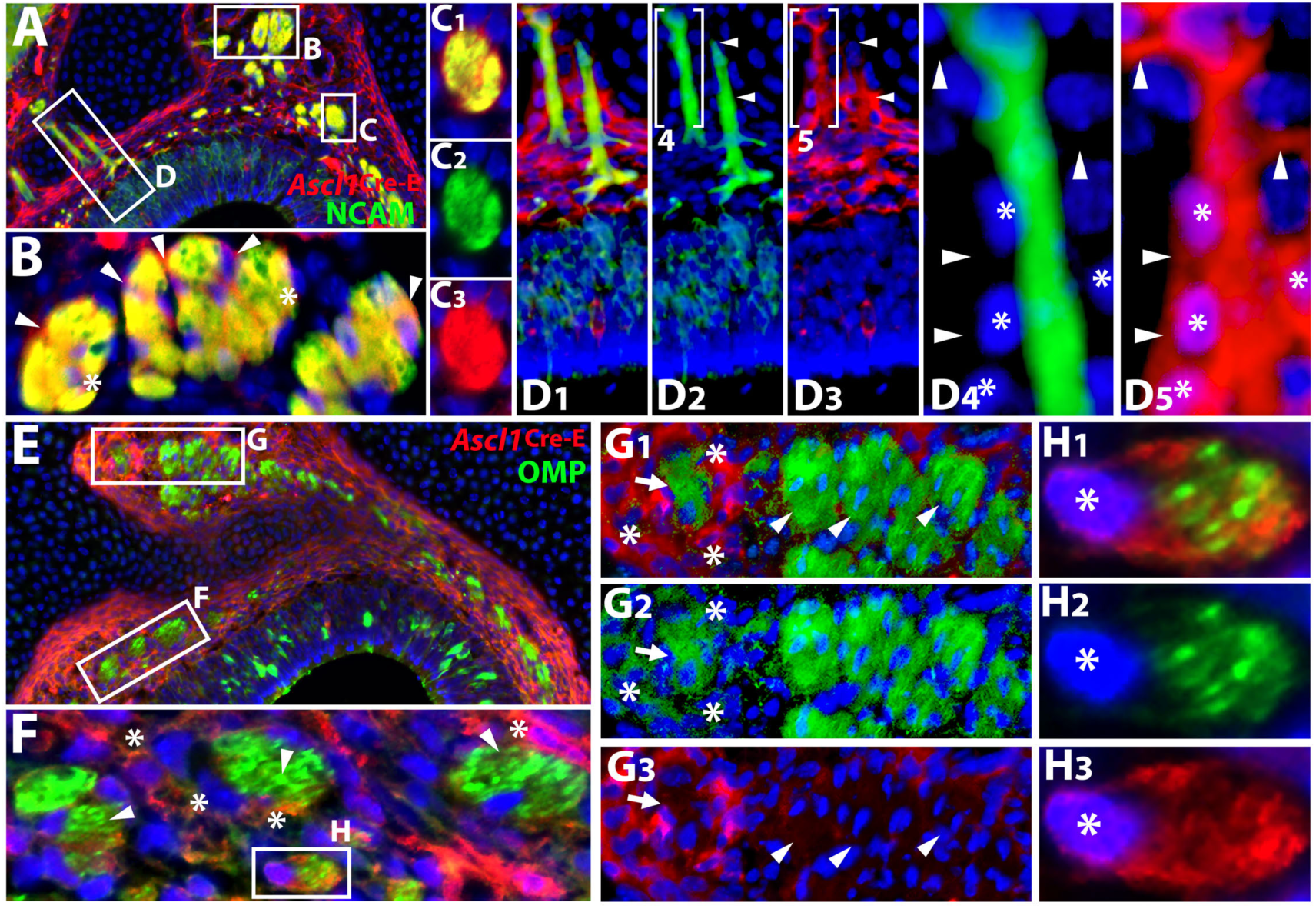
Early-specified Ascl1+ precursor-descended ensheathing cells envelop distinct classes of ORN axons in the lamina propria and coalescing olfactory nerve at E16.5. **A**) Fascicles of NCAM+ axons coalesce in the ON dorsally (boxes). **B**) Some of these fascicles have multiple NCAM+ regions bounded by septae (arrowheads) of potential E10.5-R *Ascl1*^Cre-E^+ ensheathing cells whose DAPI+ nuclei are also seen within and at the margin of these fascicles (asterisks). **C**_**1-3**_) A single large bundle of NCAM+ axons in the proximal ON enveloped by E10.5-R *Ascl1*^Cre-E^+ensheating cell processes. These processes make septa that define smaller fascicles. These E10.5-R *Ascl1*^Cre-E^+ processes may originate in distal labeled ensheathing cells since the adjacent nuclei seem to define cells that are E10.5-R *Ascl1*^Cre-E^+ -. **D1-5**) Linear-extended fascicles of NCAM+ ORN axons (e.g., bracket) grow through a more broadly distributed population of E10.5-R *Ascl1*^Cre-E^+ ensheathing cells and processes. E10.5-R *Ascl1*^Cre-E^+ cells that are not NCAM+ (asterisks, **D4**,**5**) ensheath NCAM+ axons. **E**) OMP+ E10.5-R *Ascl1*^Cre-E^-ORN axons constitute subsets of larger ON fascicles that are bounded by E10.5-R *Ascl1*^Cre-E^+ ensheathing cells and processes. **F**) Multiple OMP+ ORN axon fascicles (arrowheads) surrounded by E10.5-R *Ascl1*^Cre-E^+ ensheathing cells (asterisks). **G**_**1-3**_) Multiple OMP+ ORN axon fascicles (arrowheads) do not have adjacent enveloping E10.5-R *Ascl1*^Cre-E^+ ensheathing cells. In contrast, some OMP+ ORN axon fascicles (arrow) are surrounded by an aggregate of E10.5-R *Ascl1*^Cre-E^+ presumed ensheathing cells (asterisks). **H**_**1-3**_) A single E10.5-R *Ascl1*^Cre-E^+ ensheathing cell with a DAPI+ nucleus (left) envelops multiple small OMP+ ORN axon fascicles. The cytoplasm of the E10.5-R *Ascl1*^Cre-E^+ ensheathing cell is fenestrated at the sites where the OMP+ ORN fascicles are found.

The relationship between E10.5-R *Ascl1*^Cre-E^+ ensheathing cells and OMP+ axons at E16.5 is distinct from that with NCAM+ axons (**Figure 7E-G**). When visualized with OMP+ axons, several ensheathing cell characteristics of the *Ascl1*^Cre-E^+ cells can be resolved. The *Ascl1*^Cre-E^+ cells and processes are not labeled for OMP (**Figure 7E**). Instead, these cells form a capsule that completely envelops multiple OMP+ and presumed OMP-axon fascicles as they exit the OE into the lamina propria (**Figure 7E**). In most axon fascicles within the lamina propria, OMP+ ORN axons are intercalated with presumed OMP-axons (**Figure 7F**; *arrowheads*). The registration of partially OMP+ fascicles and E10.5-R *Ascl1*^Cre-E^+ cells is incomplete (**Figure 7G**_**1-3**;_ compare asterisks to arrowheads/arrow). Single *Ascl1*^Cre-E^+ ensheathing cells define subsets of OMP+ axon fascicles. Lamellar extensions of single *Ascl1*^Cre-E^+ cells invest small bundles of OMP+ axons (**Figure 7H**_**1-3**_). It is likely that ensheathing cells that invest additional OMP+ fascicles may either be derived from a non-Ascl+ precursor lineage or an Ascl1+ lineage that is temporally distinct from the E10.5-R cohort visualized here.

## DISCUSSION

Early OE precursors integrate temporal distinctions, molecular identity, position and signals to establish spatially restricted lineage cohorts with selective fates during initial olfactory pathway development. A temporally distinct Ascl1+ precursor-derived dorsomedial cohort, apparently descended from Pax7+/Meis1+ dorsolateral progenitors, is established during early OE differentiation (**Figure 8**). These Ascl1+ precursor-derived cells are more proliferative than Pax7+/Meis1+ precursors, respond selectively to local signals, and are more likely to generate migratory mass cells than ORNs during initial OE differentiation. Parallel analysis of modes of division of medial versus lateral OE precursors with defined molecular identities indicates that medial versus lateral Six1+, Pax7+, Ascl1+, and neurogenic precursors vary in their capacity to proliferate and divide symmetrically versus asymmetrically. These temporally distinct Ascl1+ precursors generate regionally restricted subsets of ORNs, OE supporting cells, precursors, and ensheathing cells with distinct relationships with actively growing versus mature ORN axons. This temporal, molecular, and spatial precursor diversity may establish a template in the nascent OE for cellular or regional diversity in the mature OE.

**Figure 8:**
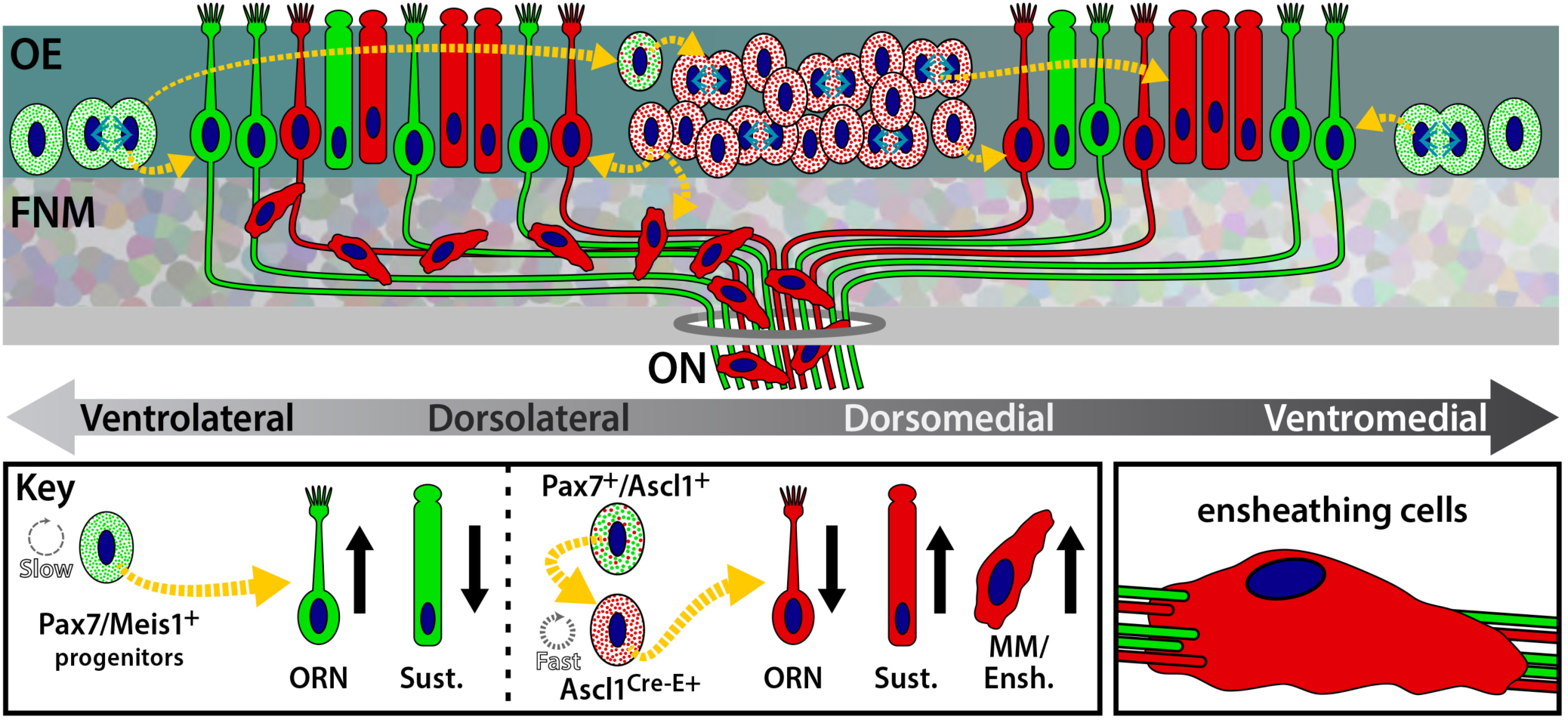
A schematic of the contribution of a temporally defined, molecularly distinct Ascl1+ derived early precursor cohort and their progeny during the initial differentiation of the olfactory epithelium (OE) and olfactory nerve (ON). Our data suggests that Meis1+/Pax7+ slowly dividing progenitors give rise to a small number of Pax7+descended/Ascl1+ precursors that generate a larger number of precursors that variably express Ascl1, are more rapidly proliferating. These precursors give rise to substantial populations of migratory mass/ensheathing cells that translocate to the frontonasal mesenchyme (FNM), associated with growing olfactory receptor neuron (ORN) axons (see inset, lower right), sustentacular cells, and a more modest population of mostly mature (i.e. NCAM-/OMP+) ORNs.

### Conditional recombination as a tool to trace embryonic molecular/temporal OE lineages

We used conditional recombination of a reporter to identify temporally distinct, lineally-related subsets of Ascl1+ OE precursors (Cau et al., 2002; Cau et al., 1997; Guillemot et al., 1993; Murray et al., 2003; Tucker et al., 2010) shortly after the initial OE morphogenesis. Similar approaches have been used to assess temporally discrete populations of Ascl1+ precursors in the spinal cord, cerebellum, and cerebral cortex (Allaway et al., 2020; Battiste et al., 2007; Sudarov et al., 2011; Vue et al., 2014) as well as for Isl1+ OE precursor fates (Taroc et al., 2020). The central role suggested for Ascl1 and other neurogenic genes in olfactory pathway development (Cau et al., 2002; Guillemot et al., 1993; Murray et al., 2003) led us to these experiments focused on Ascl1+ OE cells at the earliest stages of olfactory pathway differentiation. Conditional fate mapping of adult OE progenitors has also been used to evaluate ongoing OE neurogenesis or regeneration (Leung et al., 2007; Packard et al., 2011). These studies, however, have not focused upon precursor identity over short intervals. Conditional recombination in early (E10.5) Ascl1+ precursors assessed after two relatively short survival intervals— E11.5 and E16.5—provides a dynamic view of distinct lineage cohorts. We attempted the same conditional recombination approach using a *Pax7*^Cre-ERT^ (*Pax7*^Cre-E^) at E10.5. Extensive recombination was seen in Pax7+ lateral fnm, as expected based upon localization of Pax7 protein and mRNA (LaMantia et al., 2000; Stoykova and Gruss, 1994). We did not, however, identify any *Pax7*^Cre-E^+ OE cells at E11.5 or E16.5 in multiple experiments using two distinct *Pax7*^Cre-E^ lines (Lepper et al., 2009; Lepper and Fan, 2010; Murphy et al., 2011). This may reflect incomplete Cre expression from the *Pax7* locus in distinct cell classes or low recombination probability in slowly proliferating Pax7+/Meis1+ early OE progenitors (Tucker et al., 2010). Apparently, conditional recombination-mediated lineage mapping depends upon efficiency as well as kinetics and location of precursors that express genes encoded by loci into which Cre drivers have been inserted

### Molecular versus temporal OE precursor diversity

OE precursor specification via localized signals from the OE or fnm drives initial olfactory pathway development (Anchan et al., 1997; Balmer and LaMantia, 2004, 2005; LaMantia et al., 2000; LaMantia et al., 1993; Murray et al., 2003), mature OE regeneration and function (reviewed by Dibattista et al., 2020; Schwob et al., 2017), and genesis of olfactory ensheathing cells that support axon growth (reviewed by Franssen et al., 2007; Gomez et al., 2018). Most developmental or adult analyses define precursor diversity primarily by expression of molecular determinants—particularly bHLH transcription factors like Ascl1, Ngn2, Hes1, and NeuroD (reviewed by Kam et al., 2014)—or proliferative state. Less attention has been paid to temporal diversity of nascent or mature OE precursors or discontinuities in their distribution or fates. The E10.5-R *Ascl1*^Cre-E^+ precursor population we identified would not be recognized based only upon molecular marker expression or proliferative state. Moreover, we did not find a subset of cells for any of the additional transcription factors we analyzed (Six1, Meis1, Pax7) that was diagnostic for this temporally defined precursor population. Time, in this instance, has substantial consequences for otherwise indistinguishable precursors. This temporal identity is likely matched by a diagnostic molecular signature; however, that signature remains to be identified. Increased proliferation, limited neurogenesis, selective response to local signals and different modes of division we identified may determine or reflect temporal aspects of OE precursor identity. Our *in vivo* conditional recombination experiments and *in vitro* precursor division analysis indicate that all of these characteristics distinguish subsets of OE precursors. The subsequent fates of these precursors suggest that concatenated temporal specification of molecularly similar OE cells may establish a spatially restricted mosaic of OE progeny.

### Position and time define early OE precursor Identity

Our results suggest a novel role for an established inductive signaling mechanism that drives olfactory placode specification in parallel with that at other sites of non-axial mesenchymal/epithelial induction (reviewed by LaMantia, 2020). OE morphogenesis and ON differentiation reflects local signals from neural crest-derived mesenchyme or adjacent cranial ectoderm (Bhasin et al., 2003; Forni et al., 2013; LaMantia et al., 2000; Szabo-Rogers et al., 2009) as well as signals from the OE itself (Gokoffski et al., 2011) and amniotic fluid (AF) at the apical OE surface (Chau et al., 2015). Thus, at any time during OE morphogenesis, any single OE precursor is likely to be influenced by a positionally distinct, concentration-dependent matrix of signals that either establish or constrain subsequent proliferative or differentiation capacity. Growth of the frontonasal process likely changes this signaling matrix over time, continuously “updating” identities in temporal/spatial lineage cohorts. Spatial restriction of the *Ascl1*^Cre-E^+ OE transcriptional lineage cohort, as well as enhanced expression of pSMAD, suggests that these cells may be specified within a signaling matrix established by the OE, neural crest-derived fnm and AF, all of which provide TGFβ signals by E10.5 (LaMantia et al., 2000; Maier et al., 2010; Panaliappan et al., 2018).

The localization of Bmps, especially Bmp4, from fnm and cranial ectoderm (LaMantia et al., 2000; Shou et al., 2000); Bmp4’s differential effects on OE neurogenesis (LaMantia et al., 2000; Maier et al., 2010; Panaliappan et al., 2018; Zhu et al., 2016) and our current results showing enhanced frequency of pSMAD+ cells in the dorsomedial *Ascl1*^Cre-E^+ precursor domain suggest that Bmp4, additional Bmp family members, or other Tgfβ signals influence the *Ascl1*^Cre-E^+ OE precursor cohort. Additional signals, including Retinoic Acid, Sonic hedgehog, Fgf8, Wnts and Notch may also influence regional restriction of temporally and spatially distinct cohorts in other OE domains (Balmer and LaMantia, 2004; Herrick et al., 2018; Lassiter et al., 2014; Rawson and LaMantia, 2007; Tucker et al., 2010; Whitesides et al., 1998).

Recent work shows that subsets of Fgf20+ OE precursors, further regulated by Wnt signals guide OE turbinate morphogenesis (Yang et al., 2018). Thus, serial signaling that integrates position and time may underlie transient specification of OE lineage cohorts to influence subsequent OE development, including the E10.5-R *Ascl1*^Cre-E^+ population we identified.

### The relationship of OE precursor transcriptional and temporal/spatial identity

The *Ascl1*^Cre-E^+ temporal precursor cohort has distinct relationships with OE cells that express several additional transcription factors that are essential for optimal OE neurogenesis. Six1 (Chen et al., 2009; Ikeda et al., 2010; Ikeda et al., 2007), Meis1 (Parrilla et al., 2016; Tucker et al., 2010), and Sox2 (Dvorakova et al., 2020; Packard et al., 2011; Panaliappan et al., 2018; Tucker et al., 2010) have been implicated in regulation of OE precursor lineage progression and fate. It is not clear, however, whether OE precursors expressing any of these transcriptional regulators have consistent, singular properties.

Despite shared Sox2 expression, proliferation and modes of division of Six1+, Pax7+, or Ascl1+ OE precursors differ based upon medial versus lateral origin. These distinctions may reflect local signals (see above) that impose positional identity on OE precursor subpopulations. Alternately, they may reflect local cell-cell interactions that nevertheless depend upon the influence of immediate neighbors of similar or distinct identities (Karpinski et al., 2021). Temporal specification of subsets of precursors may provide greater diversity of potential interactions. Apparently, OE precursor proliferative capacity and mode of division is established by integrating molecular expression signatures, signaling history, position, and temporal relationships among OE precursors.

### Temporally distinct OE precursor cohorts, cell fate, and ORN diversity

Temporally distinct cohorts of OE precursors with distinct molecular identity and axial locations may contribute to the regional or “zonal” organization of ORNs distinguished by monoallelic odorant receptor expression and segregated projections to the OB in the mature mammalian olfactory pathway (Buck, 1996; Mori et al., 2000; Ressler et al., 1993; Vassar et al., 1993; Zapiec and Mombaerts, 2020).

E10.5-R *Ascl1*^Cre-E^+ precursor progeny identified at E16.5 includes a subset of ORNs limited to an OE region that corresponds approximately to Zone 1 (reviewd by Mori et al., 2000) or Zolf24/Zolf16 defined more recently (Zapiec and Mombaerts, 2020), based upon monoallelic ORN expression of odorant receptor genes. Some Ascl1 lineage-derived ORNs in this subregion at E16.5 appear more mature based upon co-expression of OMP (Graziadei et al., 1980; Verhaagen et al., 1989) and exclusion of NCAM (Miragall et al., 1989; Yamashita et al., 1998). It is possible that the E10.5-R *Ascl1*^Cre-E^+ lineage cohort generates a “pioneer” population of Zone 1 ORNs whose axons establish distinct OB projections. We detect few BrdU+/*Ascl1*^Cre-E^+ presumed OE resident progenitors at E16.5. The E10.5-R *Ascl1*^Cre-E^+ ORNs may establish initial restricted projections which are replaced by ORNs generated by later arising precursors. Whether ORNs generated by temporally distinct *Ascl1*^Cre-E^+ precursors endure beyond fetal life, or small number of precursors from this cohort expand clonally to facilitate genesis of distinct subsets of ORNs during subsequent OE maturation or ongoing regeneration remains to be determined. A more detailed understanding of diversity of OE precursor lineages, their early derivations, and subsequent fates as either biased precursors for specific OE cell classes or multipotent OE progenitors is clearly necessary to better predict consequences of OE pathogenesis, including viral infection and damage (Cooper et al., 2020), and to optimize OE regeneration and olfactory functional recovery.

### A distinct lineage of olfactory ensheathing cells coordinates ORN axon growth

The earliest cohort of migratory mass cells—forerunners of olfactory ensheathing cells— and subsequently a subset of ensheathing cells themselves are derived from the *Ascl1*^Cre-E^+ OE lineage cohort (**Figure 8**). *Ascl1*^Cre-E^+ migratory mass cells are unlikely to include GnRH neurons, which are generated despite full loss of Ascl1 function (Tucker et al., 2010). Indeed, presumed GnRH progenitors in the ventromedial OE express nestin, a general neural progenitor marker, but not Ascl1 (Kramer and Wray, 2000). Our Pax7 protein localization and Cre-dependent lineage tracing suggest that GnRH neurons may be derived from Pax7+ ventromedial OE precursors. The apparent absence of Ascl1 expression from the fnm (Guillemot and Joyner, 1993), which is largely derived from neural crest (Karpinski et al., 2016; Osumi-Yamashita et al., 1994; Serbedzija et al., 1992) and the labeling of a substantial number of these cells at E11.5 via Ascl1-mediated recombination in the OE make it unlikely that migratory mass or olfactory ensheathing cells are generated solely by fnm neural crest (Barraud et al., 2010; Forni et al., 2011; Katoh et al., 2011). Instead, our current *in vivo* data are consistent with earlier *in vitro* observations that at least the initial population of migratory mass cells is generated primarily from olfactory placodal ectoderm (LaMantia et al, 2000; Rawson et al, 2010). Substantial populations of olfactory ensheathing cells at E16.5 descend from E10.5-R *Ascl1*^Cre-E^+ mediodorsal OE precursors. These ensheathing cells have distinct relationships with subsets of growing (NCAM+) as well as more mature (OMP+) ORN axons. This suggests that ensheathing cell identities may be restricted in register with lineage-related derivation of ORNs. Accordingly, both the supporting glia and projecting ORNs may be derived from common temporally and spatially defined OE precursors. Such a relationship, in concert with other developmental and activity-dependent mechanisms, might facilitate subsequent high-fidelity regeneration of ORN axons (Gogos et al., 2000) based upon lineage-dependent matched molecular affinities.

## ACKNOWLEDGEMENTS

This work was supported by NIDCD DC011534 and NICHD HD083157 to A-S. L. We thank Anastas Popratiloff and the staff of the GW Nanofabrication and Imaging Center for assistance with confocal microscopic imaging. We thank Mike Fox for his thoughtful comments on the manuscript, and Connor Siggins for reviewing the final version.

## Notes

### Competing Interest Statement

The authors have declared no competing interest.

